# Blocking HXA_3_-mediated neutrophil elastase release during *S. pneumoniae* lung infection limits pulmonary epithelial barrier disruption and bacteremia

**DOI:** 10.1101/2024.06.25.600637

**Authors:** Shuying Xu, Shumin Tan, Patricia Romanos, Jennifer L. Reedy, Yihan Zhang, Michael K. Mansour, Jatin M. Vyas, Joan Mecsas, Hongmei Mou, John M. Leong

## Abstract

*Streptococcus pneumoniae* (*Sp*), a leading cause of community-acquired pneumonia, can spread from the lung into the bloodstream to cause septicemia and meningitis, with a concomitant three-fold increase in mortality. Limitations in vaccine efficacy and a rise in antimicrobial resistance have spurred searches for host-directed therapies that target pathogenic immune processes. Polymorphonuclear leukocytes (PMNs) are essential for infection control but can also promote tissue damage and pathogen spread. The major *Sp* virulence factor, pneumolysin (PLY), triggers acute inflammation by stimulating the 12-lipoxygenase (12-LOX) eicosanoid synthesis pathway in epithelial cells. This pathway is required for systemic spread in a mouse pneumonia model and produces a number of bioactive lipids, including hepoxilin A3 (HXA_3_), a hydroxy epoxide PMN chemoattractant that has been hypothesized to facilitate breach of mucosal barriers. To understand how 12-LOX-dependent inflammation promotes dissemination during *Sp* lung infection and dissemination, we utilized bronchial stem cell-derived air-liquid interface (ALI) cultures that lack this enzyme to show that HXA_3_ methyl ester (HXA_3_-ME) is sufficient to promote basolateral-to-apical PMN transmigration, monolayer disruption, and concomitant *Sp* barrier breach. In contrast, PMN transmigration in response to the non-eicosanoid chemoattractant fMLP did not lead to epithelial disruption or bacterial translocation. Correspondingly, HXA_3_-ME but not fMLP increased release of neutrophil elastase (NE) from *Sp*-infected PMNs. Pharmacologic blockade of NE secretion or activity diminished epithelial barrier disruption and bacteremia after pulmonary challenge of mice. Thus, HXA_3_ promotes barrier disrupting PMN transmigration and NE release, pathological events that can be targeted to curtail systemic disease following pneumococcal pneumonia.

**Importance:** *Streptococcus pneumoniae* (*Sp*), a leading cause of pneumonia, can spread from the lung into the bloodstream to cause systemic disease. Limitations in vaccine efficacy and a rise in antimicrobial resistance have spurred searches for host-directed therapies that limit pathologic host immune responses to *Sp*. Excessive polymorphonuclear leukocyte (PMN) infiltration into *Sp*-infected airways promotes systemic disease. Using stem cell-derived respiratory cultures that reflect *bona fide* lung epithelium, we identified the eicosanoid hepoxilin A3 as a critical pulmonary PMN chemoattractant that is sufficient to drive PMN-mediated epithelial damage by inducing the release of neutrophil elastase. Inhibition of the release or activity of this protease in mice limited epithelial barrier disruption and bacterial dissemination, suggesting a new host-directed treatment for *Sp* lung infection.

## Introduction

*Streptococcus pneumoniae* (*Sp*; also known as the pneumococcus) is a Gram-positive bacterium that asymptomatically colonizes the nasopharynx of 5-10% of healthy adults, but can spread to the lower respiratory tract and is the most frequent cause of community-acquired pneumonia (1). Subsequent bacterial translocation from the airway into the bloodstream can lead to invasive disease, such as septicemia and meningitis, events associated with a three-fold increase in mortality (2). Invasive pneumococcal infections result in approximately 14 million cases and one million deaths annually worldwide (3). Vaccination and antimicrobials are first-line strategies in combating pneumococcal diseases. However, the rapid rise of antibiotic resistance and the limited antigenic breadth of effective vaccines have fueled interest in treatment strategies that focus on diminishing tissue-destructive host immune responses (4–7).

Pneumococcal infection of lung mucosa drives robust recruitment of polymorphonuclear leukocytes (PMNs, or neutrophils), leading to the acute inflammation that is a hallmark of this infection (1). PMNs confront invading *Sp* with multiple antibacterial mechanisms, including release of reactive oxygen species (ROS) (8), neutrophil extracellular traps (NET) (9), and/or proteases such as cathepsin G (CG) and neutrophil elastase (NE) (10). Indeed, neutropenic individuals or neutrophil-depleted mice are highly susceptible to systemic *Sp* infection (11, 12). Nevertheless, sustained pulmonary accumulation of PMNs increases airway permeability with a concomitant risk of disseminated infection (13, 14). Protease inhibitors that diminish PMN infiltration also reduce bacteremia and lethality after *Sp* pulmonary challenge of mice (15, 16). Finally, mice that retain high numbers of pulmonary PMNs suffer higher levels of bacteremia and mortality (17–20), and depletion of PMNs 18 hours post-infection (h.p.i.) mitigates disease and pathogen spread (21).

Chemotactic cues not only recruit PMNs but also influence their tissue-destructive character (22–24). Hence, in addition to their recruitment, PMN-directed pathologies may result from enhanced tissue-damaging PMN activities (24). The major *Sp* virulence factor pneumolysin (PLY), a cytolysin that drives tissue damage and promotes early bacteremia (25–27), stimulates the 12-lipoxygenase (12-LOX) pathway in epithelial cells and results in the synthesis and apical secretion of eicosanoid PMN chemoattractants (17, 28, 29). Among 12-LOX-generated bioactive lipid mediators (30), the hydroxy epoxide hepoxilin A3 (HXA_3_) is a potent chemoattractant (31) that orchestrates mucosal inflammation during both intestinal (32, 33) and pulmonary infections (34). Like other chemoattractants (23, 35), HXA_3_ has both chemotactic and non-chemotactic effects on PMNs (36), triggering intracellular calcium release (36), promoting PMN survival (37), inducing NET formation (38), and stimulating the release of additional arachidonic acid metabolites (39). Notably, genetic ablation or chemical inhibition of 12-LOX drastically reduces PMN infiltration, bacteremia, and mortality following *Sp* lung challenge of mice (17, 29), suggesting that barrier disruption and systemic *Sp* disease could be mitigated by modulation of PMN effector functions that are enhanced by one or more products of the 12-LOX pathway.

The tissue-destructive functions of PMNs are dramatically altered upon exposure to bacterial factors (24, 40, 41), but the effect of HXA_3_ on PMNs in the context of *Sp* infection has not been examined. In addition, 12-LOX promotes the production of numerous bioactive lipids (30), and although HXA_3_ has been hypothesized to be the essential driver in PLY-promoted *Sp* dissemination from the lung, this eicosanoid has not been directly implicated in the *Sp*- or PLY-driven PMN chemotaxis. These limitations are in part a reflection of the instability of HXA_3_ in aqueous environments (32), as well as the lack of an easily manipulated *in vitro* experimental model that faithfully reflects *Sp*-mediated inflammation and bacterial translocation across an epithelial barrier. Indeed, the respiratory epithelial culture models previously applied to *Sp* infection are typically based on immortalized cell lines that lack the cellular diversity and *bona fide* barrier function integral to airway epithelium (42). Here, we characterized the role of PLY in promoting PMN transmigration and epithelial compromise using air-liquid interface (ALI) monolayers derived from bronchial stem cells that recapitulate key features of the airway epithelium. Moreover, ALI monolayers genetically ablated for 12-LOX deficient permitted the demonstration that HXA_3_ methyl ester (HXA_3_-ME), a stable and active version of HXA_3_, is sufficient to promote PMN transmigration and *Sp* barrier breach. Corresponding studies of the signaling capacities of HXA_3_-ME on PMN in the context of *Sp* infection showed that HXA_3_ is not only a central driver of PMN transmigration across infected epithelium but also enhances the tissue-damaging proteolytic activity of PMNs. Targeting this HXA_3_-promoted activity mitigated systemic disease following *Sp* pulmonary challenge of mice, illustrating its therapeutic potential as a host-directed therapy for *Sp* infection.

## Results

### The 12-LOX pathway, stimulated by PLY-producing *Sp*, promotes PMN infiltration, lung permeability, and bacteremia following *Sp* lung infection in mice

Activation of the airway epithelial cell 12-LOX pathway is triggered by *Sp* pneumolysin (29). We intratracheally (*i.t*.) inoculated BALB/c mice with 1×10^7^ CFU of WT *Sp* TIGR4 or the isogenic PLY-deficient mutant *Sp* TIGR4 *Δply*. At 18 hours post infection (h.p.i.), the two strains reached similar lung burdens (Figure 1a, “WT” vs. “*Δply*”), consistent with previous reports (43, 44). Both strains also induced pulmonary inflammation, but consistent with the ability of PLY to stimulate the 12-LOX pathway and increase inflammation (17, 26), PMN pulmonary infiltration was 1.5-fold higher in mice infected with WT *Sp* compared to *Sp Δply* (p < 0.01; Figure 1b, “WT” vs. “*Δply*”). To assess damage to the lung barrier, at 18 h.p.i. we delivered 1 mg of 70 kDa FITC-dextran intravenously into infected mice and, after 30 minutes, measured the fluorescence signal in lung homogenates relative to that of serum. Infection by WT *Sp* increased lung permeability more than two-fold relative to uninfected mice (p < 0.01), whereas infection with *Sp Δply* had no effect (Figure 1c, “WT” vs. “*Δply*”). Mirroring the increased lung permeability to FITC-dextran, WT *Sp* infection resulted in a ten-fold higher level of bacteremia compared to *Sp Δply* infection (p < 0.05; Figure 1d, “WT” vs. “*Δply*”).

**Figure 1.**
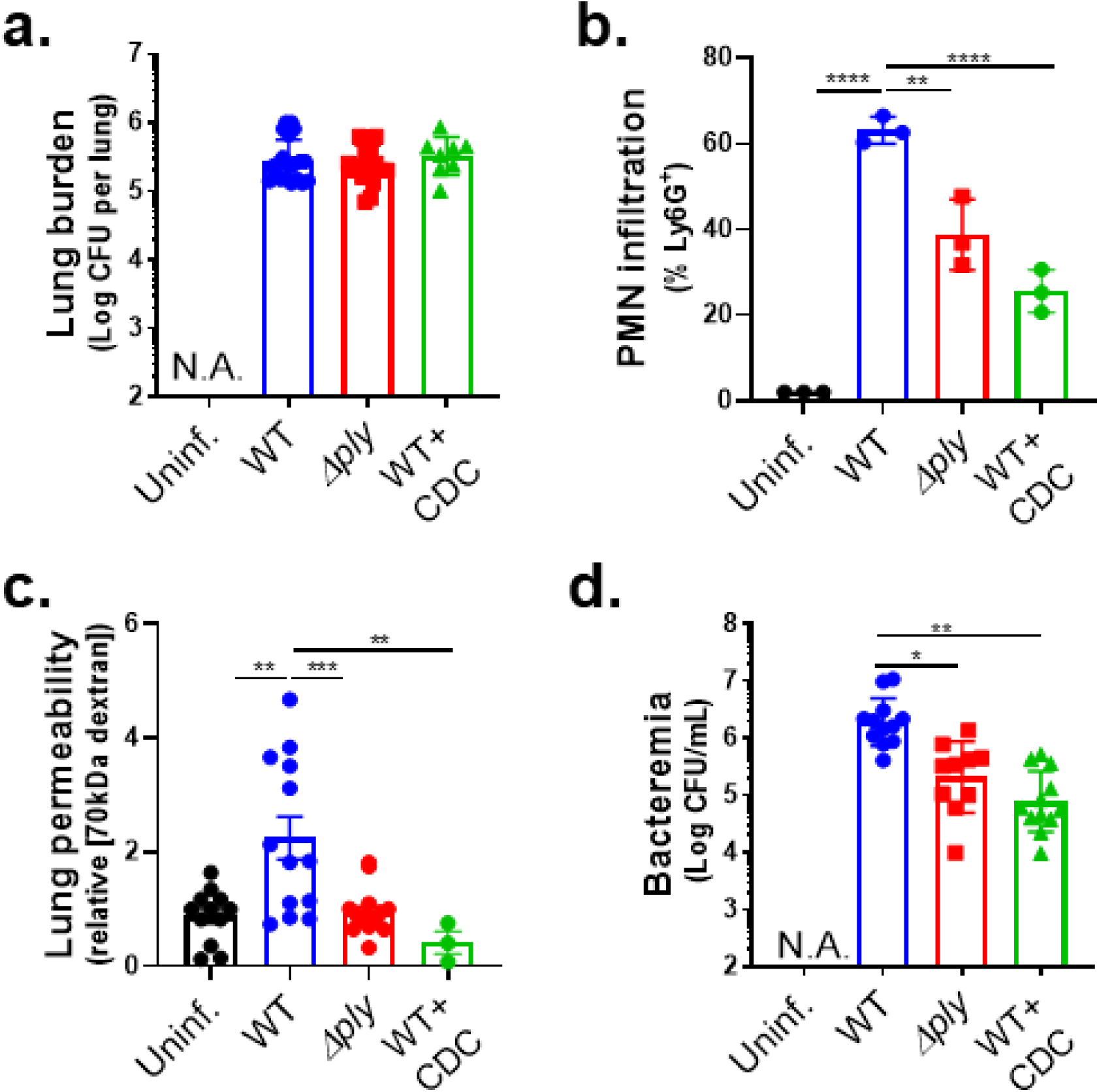
The 12-LOX pathway, stimulated by PLY-producing *Sp*, promotes PMN infiltration, lung permeability and bacteremia following *Sp* lung infection in mice. BALB/c mice were infected *i.t.* with 1 × 10^7^ CFU wild type (WT) or PLY-deficient mutant (*Δply*) TIGR4 *Sp* for 18 h, with or without *i.p* injection of 8 mg/kg of the 12-LOX inhibitor CDC. **(a)** Bacterial lung burden determined by measuring CFU in lung homogenates. **(b)** PMN infiltration determined by flow cytometric enumeration of Ly6G^+^. **(c)** Lung permeability quantitated by measuring the concentration of 70 kDa FITC-dextran in the lung relative to serum after *i.v.* administration. **(d)** Bacteremia measured by enumerating CFU in serum. Each panel is representative of three independent experiments, or pooled data from three independent experiments. Error bars represent mean ± SEM. Statistical analysis was performed using ordinary one-way ANOVA: *p-value < 0.05, **p-value < 0.01, ***p-value < 0.001, ****p-value < 0.0001.

We previously found that inhibition of 12-LOX activity by *i.p* injection of cinnamyl-3,4-dihydroxy- α-cyanocinnamate (CDC) did not affect Sp lung burden but curtailed PMN lung infiltration in C57BL/6 (B6) mice (17). Here, after infection of BALB/c mice with WT *Sp*, CDC treatment similarly diminished lung PMN infiltration (p < 0.0001) without altering lung burden (Figure 1a-b, “WT + CDC”). Lung barrier disruption and *Sp* dissemination also depended on 12-LOX activity because CDC treatment of *Sp*-infected mice resulted in lower FITC-dextran leakage (p < 0.01; Figure 1c) and bacteremia at 18 h.p.i. (p < 0.01; Figure 1d). Therefore, 12-LOX activation by PLY promoted PMN infiltration to the lungs, an event that correlated with increased lung permeability and *Sp* spread to the bloodstream.

### The 12-LOX pathway promotes PMN transmigration and epithelial barrier breach upon apical infection of ALI monolayers by PLY-producing *Sp*

Recent advances in airway stem cell biology have allowed for the generation of genetically tractable *in vitro* stem cell-derived epithelial cultures with organized architecture and functional attributes of the airway mucosa, including beating cilia, apical mucus production, and a robust junctional barrier (45). To identify key steps underlying the promotion of bacteremia by PLY and 12-LOX activation, we modeled interactions between *Sp* and PMNs at the airway epithelium by culturing human airway basal stem cells (BSCs) on 3 µm pore size Transwell filters. After growth to confluency, media was removed from the apical side of the monolayers, a step that triggers the differentiation of the stem cells to form a monolayer containing the diverse airway epithelial cell types (42), including ciliated cells, mucus-producing goblet cells, and secretory club cells, found in *bona fide* airway epithelium. We then added 1×10^6^ PMNs isolated from human peripheral blood to the basolateral surface of these air-liquid interface (ALI) cultures and assessed their movement to the apical side upon *Sp* infection.

Two hours of apical infection with 1×10^7^ *Sp*/Transwell induced robust PLY-dependent PMN transmigration across human ALI monolayers, with WT *Sp* triggering two-fold greater migration compared to *Sp Δply* (p < 0.0001; Figure 2a, “Human ALI”). WT *Sp* infection of monolayers pre-treated with CDC failed to trigger PMN transmigration (Figure 1a, “WT+CDC”), suggesting that PMN transmigration across *Sp*-infected ALI monolayers was dependent on eicosanoid lipid mediators produced by 12-LOX, recapitulating our findings during pulmonary *Sp* challenge in mice.

**Figure 2.**
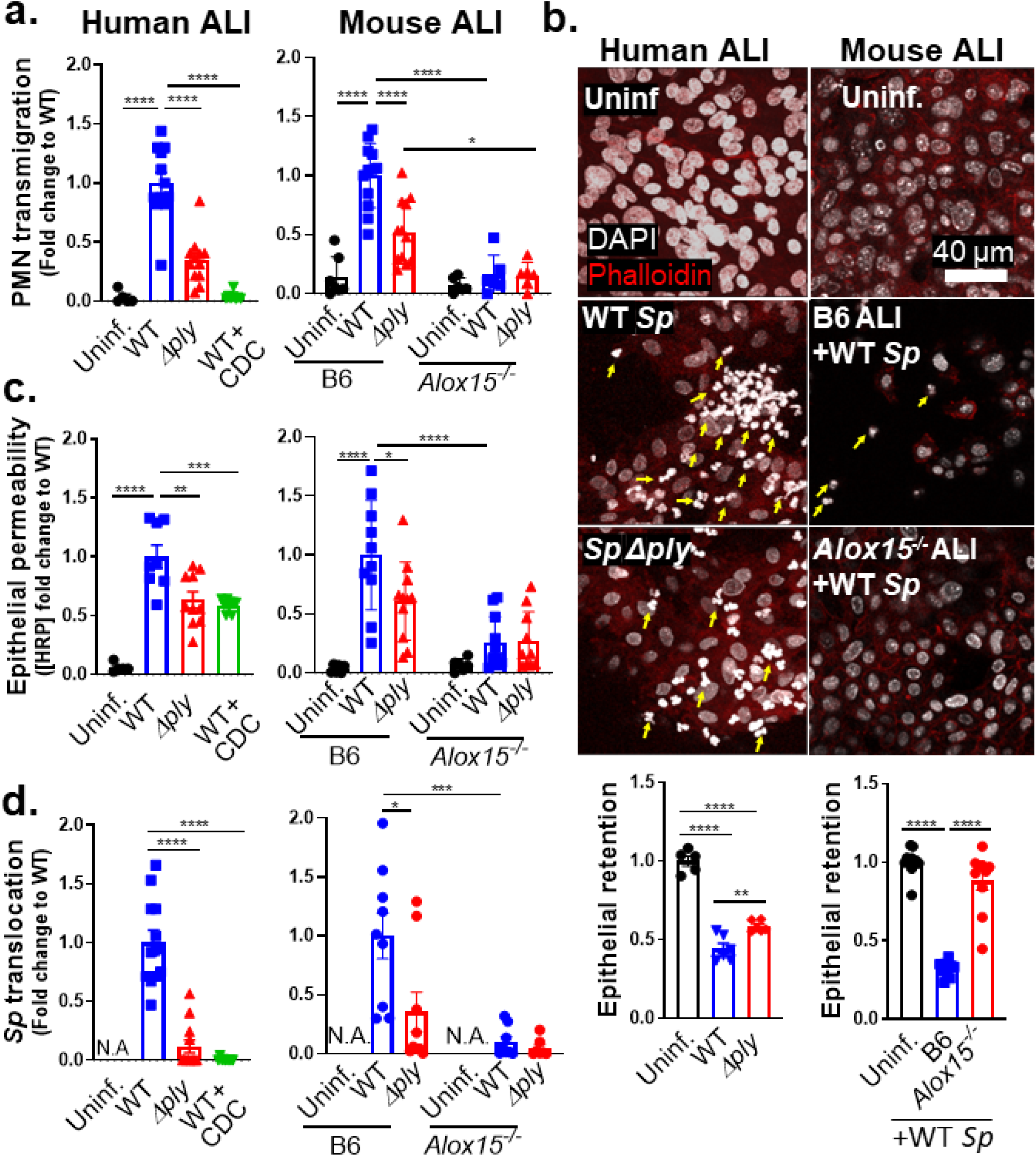
The 12-LOX pathway promotes PMN transmigration and epithelial barrier breach upon apical infection of ALI monolayers by PLY-producing *Sp*. Human BSC-derived ALI monolayers (left column) or WT B6 and 12-LOX-deficient *Alox15*^-/-^ mouse BSC-derived ALI monolayers (right column) were apically infected with 1 × 10^7^ WT or *Δply Sp* in the presence of basolateral PMNs. **(a)** After 2 hours of PMN migration, the degree of transmigration as determined by MPO activity in the apical chamber. **(b)** PMN infiltration and monolayer integrity assessed by fluorescence confocal microscopy after staining nuclei with DAPI and F-actin with fluorescent phalloidin. For clarity, images shown are of extended projections (all z-sections collapsed into 1 plane). Arrows indicate examples of PMN nuclei. Scale bar = 40 μm for all images. Quantitation of epithelial retention is shown in the graph below the images, performed by enumerating epithelial cell nuclei relative to uninfected ALI in five images per experiment. **(c)** Epithelial permeability measured by HRP flux relative to monolayers infected with WT *Sp*. (d) *Sp* translocation quantitated by measuring basolateral CFU. Each panel is representative of three independent experiments, or pooled data from three independent experiments. Error bars represent mean ± SEM. Statistical analysis was performed using ordinary one-way ANOVA: *p-value < 0.05, **p-value < 0.01, ***p-value < 0.001, ****p-value < 0.0001.

Given the correlation between PMN infiltration and barrier disruption *in vivo*, we visualized monolayers by fluorescence confocal microscopy. PMNs were distinguished from ALI cells by staining cell nuclei with DAPI and visualizing their F-actin with fluorescent phalloidin. Upon infection with WT *Sp*, PMNs, identified by their multi-lobed nuclei, were found to infiltrate the epithelial monolayers in great numbers. Infection with *Sp Δply* resulted in reduced but detectable PMN infiltration (Figure 2b, “Human ALI”, yellow arrows). On the other hand, epithelial cells were lost from the Transwell filters post-PMN transmigration. To quantitate epithelial cell loss, we optimized a CellProfiler pipeline to distinguish epithelial cells from PMNs based on the size and shape of their nuclei (see Methods). Quantitation of each cell type indicated that infection with WT *Sp* and concomitant PMN migration triggered a 64% loss in epithelial cells from the monolayer (Figure 2b, “WT *Sp*”). This loss was entirely dependent on the presence of PMNs (Figures S1a-b). It was also partially dependent on PLY, because infection with *Sp Δply* resulted in a 41% (and significantly lower) loss of epithelial cells (p < 0.01; Figure 2b, “*Sp Δply*”).

To quantitate epithelial barrier function, we measured leakage of the basally loaded tracer protein HRP into the apical chamber. A 17-fold increase in HRP flux was observed after PMN transmigration induced by apical infection of ALI monolayers by WT *Sp* (p < 0.0001; Figure 2c, “Human ALI”). This level of leakage was 1.5-fold higher compared to monolayers that had been pre-treated with CDC or monolayers that were infected with *Sp Δply* (p < 0.01; Figure 2c, “Human ALI”). The diminished HRP leakage observed in the latter conditions correlated with a 25- or 9-fold decrease in cross-monolayer bacterial movement (p < 0.0001; Figure 2d, “Human ALI”). As predicted, disruption to barrier integrity depended entirely on the presence of PMNs (Fig S1c-d).

We then tested the effect of genetic ablation of 12-LOX by generating ALI monolayers from WT or 12-LOX-deficient *Alox15*^-/-^ mice (Figure 2a, “Mouse ALI”). Infection of ALI monolayers from B6 mice with WT *Sp* induced PMN transmigration 7-fold higher than basal (uninfected) levels (p < 0.0001) and 2-fold higher (p < 0.0001) than that induced by *Sp Δply* (Figure 2a, “B6”). In contrast, *Alox15*^-/-^ ALI monolayers failed to trigger significant PMN transmigration during infection by either WT or PLY-deficient *Sp* (Figure 2a, “*Alox15*^-/-^”).

Confocal microscopy analysis of monolayers after PMN migration revealed that WT infection was associated with a 68% loss of B6 ALI monolayer compared to a 12% loss of *Alox15*^-/-^ ALI monolayers (p < 0.0001; Figure 2b, “Mouse ALI”). Correspondingly, a 30-fold increase in HRP flux was detected across monolayers infected with WT *Sp* compared to uninfected monolayers (p < 0.0001; Figure 2c, “Mouse ALI”). This increase in HRP flux was promoted by both PLY and 12-LOX, because (a) *Δply Sp* infection of WT B6 monolayers resulted in 2-fold lower flux (p < 0.05); and (b) WT *Sp* infection of *Alox15*^-/-^ ALI monolayers resulted in 4-fold lower flux (p < 0.0001; Figure 2c, “Mouse ALI”).

The PLY- and 12-LOX-dependent barrier disruption correlated with enhanced *Sp* translocation across ALI monolayers, as WT *Sp* translocation across B6 monolayers was 3-fold higher than that of *Sp Δply* (p < 0.05) and 10-fold higher than that of WT *Sp* across *Alox15*^-/-^ monolayers (p < 0.001; Figure 2d, “Mouse ALI”). Notably, although PLY has diverse effects on mammalian cells (46, 47), upon infection of 12-LOX-deficient ALI, the presence or absence of PLY had no effect on barrier disruption and bacterial translocation. Thus, not only is 12-LOX-dependent PMN transmigration required for barrier breach during *Sp* infection of ALI monolayers, but the critical role of PLY in this process is the induction of the 12-LOX pathway.

### A soluble factor produced by ALI monolayers via the 12-LOX pathway upon apical *Sp* infection promotes both PMN migration and barrier disruption

Infection of WT but not 12-LOX-deficient ALI monolayers by *Sp* triggered PMN migration and barrier breach (Fig. 2; “Mouse ALI”; “*Alox15^-/-^*”). To detect putative soluble factor(s) produced by infected epithelium via the 12-LOX pathway, we first collected apical supernatants from B6 ALI monolayers that had been infected with WT *Sp* (herein referred to as “WT supernatant”), or as controls, infected with *Δply Sp* (“*Δply* supernatant”) or left uninfected (“uninfected supernatant”). (We did not include these supernatants of *Alox15^-/-^* ALI cultures because these monolayers did not support PMN migration under any conditions; Fig. 2a). Detecting factors that are capable of drawing PMNs across *Sp*-infected ALI monolayers and facilitating bacterial translocation is confounded by the further production of 12-LOX-derived products by infected cells. Hence, we added these supernatants to *Alox15*^-/-^ (not WT B6) ALI monolayers that had been apically infected with WT *Sp*. The addition of WT supernatant triggered PMN transmigration across infected *Alox15*^-/-^ ALI monolayers at a 25- and 2-fold higher level than that triggered by uninfected supernatant and *Δply* supernatant, respectively (Figure 3a).

**Figure 3.**
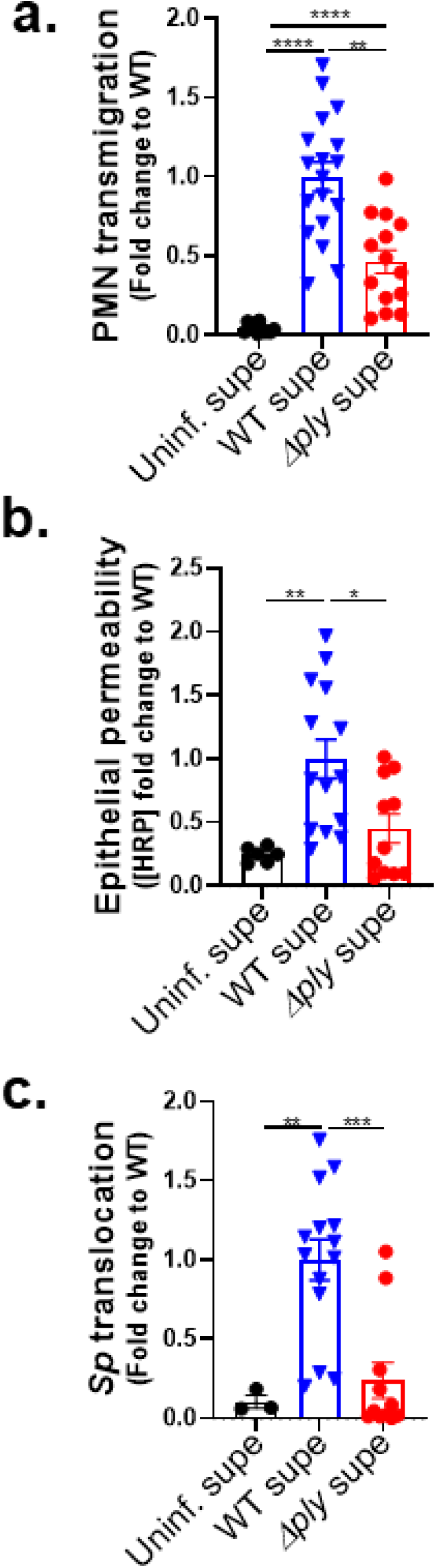
A soluble factor produced by ALI monolayers via the 12-LOX pathway upon apical *Sp* infection promotes both PMN migration and barrier disruption. *Alox15*^-/-^ mouse BSC-derived ALI monolayers were apically infected with 1 × 10^7^ WT *Sp* and transferred into apical chambers containing supernatant generated from WT *Sp* infection (WT supe) or *Δply* infection (*Δply* supe) of B6 mouse BSC-derived ALI monolayers. **(a)** After two hours of PMN migration, the degree of transmigration as determined by MPO activity in the apical chamber. **(b)** Epithelial permeability measured by HRP flux relative to monolayers infected with WT *Sp*. (c) *Sp* translocation quantitated by measuring basolateral CFU. Each panel is representative of three independent experiments, or pooled data from three independent experiments. Error bars represent mean ± SEM. Statistical analysis was performed using ordinary one-way ANOVA: *p-value < 0.05, **p-value < 0.01, ***p- value < 0.001.

To determine if PMN migration in response to a 12-LOX-dependent soluble factor (or factors) disrupted the infected monolayer, we measured cross-epithelial horseradish peroxidase (HRP) leakage. WT supernatant induced 4- and 2-fold more leakage than uninfected supernatant and *Δply* supernatant, respectively (Figure 3b). In turn, HRP leakage correlated with bacterial movement because WT supernatant was associated with 10- and 5-fold higher *Sp* translocation than uninfected and *Δply* supernatant, respectively (Figure 3c). That supernatant of epithelium infected with WT *Sp* was sufficient to rescue PMN migration across *Alox15*^-/-^ ALI monolayers, as well as concomitant barrier disruption and *Sp* translocation, affirmed the presence of a soluble mediator (or mediators) in the epithelial apical supernatant that acts as a PMN chemoattractant and drives barrier breach during *Sp* infection.

### Upon *Sp* infection of ALI monolayers, PMN transmigration induced by HXA_3_ but not fMLP promotes barrier breach

The 12-LOX pathway generates a number of bioactive lipids, but based on mucosal infection by several bacterial pathogens (32, 34, 48), hepoxilin A3 (HXA_3_) is a prime candidate for the 12-LOX-dependent chemoattractant secreted into the apical supernatant by infected B6 ALI monolayers. To test whether HXA_3_ is sufficient to trigger PMN transmigration, barrier disruption, and bacterial translocation *in vitro*, we added HXA_3_ methyl ester (HXA_3_-ME), a stable synthetic form of HXA_3,_ to the apical chamber of *Alox15*^-/-^ ALI monolayers infected with WT *Sp*, and monitored transmigration of basolateral PMNs. As controls, the well-characterized non-eicosanoid PMN chemoattractant N-formyl-L-methionyl-L-leucyl-phenylalanine (fMLP) induced PMN transmigration, whereas the HBSS buffer control did not (Figure 4a, “HBSS”, “fMLP”). We found that the apical addition of HXA_3_-ME induced PMN transmigration equivalent to that triggered by fMLP (Figure 4a, “HXA_3_”), Indicating that HXA_3_ is sufficient to induce PMN migration across *Sp*-infected ALI monolayers.

**Figure 4.**
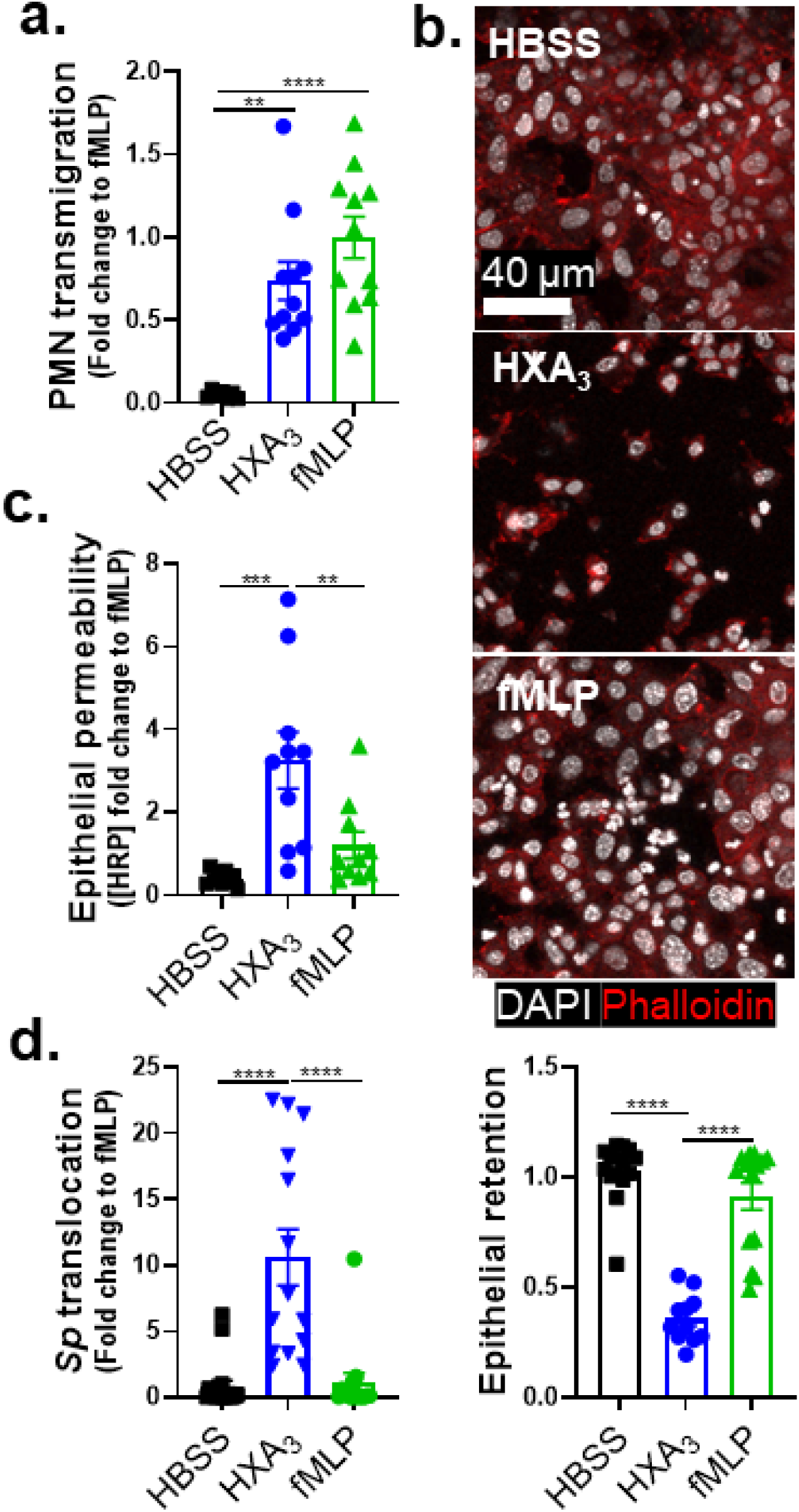
Upon *Sp* infection of ALI monolayers, PMN transmigration induced by HXA_3_ but not fMLP promotes barrier breach. *Alox15*^-/-^ mouse BSC-derived ALI monolayers were apically infected with 1 × 10^7^ WT *Sp* and transferred into apical chambers containing 10 nM HXA_3_ methyl ester (“HXA_3_”), or 10 µM fMLP, in the presence of basolateral PMNs. **(a)** After two hours of PMN migration, the degree of transmigration as determined by MPO activity in the apical chamber. **(b)** Monolayer integrity assessed by fluorescence confocal microscopy after staining nuclei with DAPI and F-actin with fluorescent phalloidin. For clarity, images shown are of extended projections (all z-sections collapsed into 1 plane). Scale bar = 40 μm for all images. Shown below the images is epithelial retention quantitated by enumerating epithelial cell nuclei relative to uninfected monolayers in five images per experiment. **(c)** Epithelial permeability measured by HRP flux relative to monolayers infected with WT *Sp*. **(d)** *Sp* translocation quantitated by measuring basolateral CFU. Each panel is representative of three independent experiments, or pooled data from three independent experiments. Error bars represent mean ± SEM. Statistical analysis was performed using ordinary one-way ANOVA: *p-value < 0.05, **p-value < 0.01, ***p-value < 0.001, ****p-value < 0.0001.

In addition to inducing PMN migration, chemoattractants can alter other PMN functional responses (24), and HXA_3_ influences a variety of PMN behaviors (36), such as intracellular calcium release (36), apoptosis inhibition (37), and NETosis (38). Indeed, despite similar levels of PMN transmigration in response to HXA_3_ and fMLP, PMN transmigration induced by fMLP was associated with retention of the epithelial monolayer integrity (Figure 4b), minimal HRP flux (Figure 4c), and the absence of *Sp* transepithelial movement (Figure 4d, “fMLP”), whereas that mediated by HXA_3_-ME induced loss of 64% of the monolayer, a 4-fold increase in HRP leakage, and a 10-fold increase in *Sp* translocation (Figure 4b-d, “HXA_3_”). These data indicate that HXA_3_ induces a mode of PMN transmigration capable of promoting barrier disruption and bacterial translocation.

### HXA_3_-stimulated PMNs generate robust NE in response to *Sp* infection

Previous studies show that PMNs respond to purified HXA_3_ by resisting apoptosis (37) and generating NETs (38), events that may reinforce PMN inflammatory potential. However, the potentially tissue-damaging state of PMNs is greatly influenced by exposure to microbial pathogens (24). Hence, to identify key features ofPMNs that may lead to monolayer disruption and *Sp* translocation, we compared the effect of HXA_3_ and fMLP on various PMN responses in the context of*Sp* infection. To begin this analysis, we first characterized PMN activities in response to *Sp* in the absence of chemoattractant. After 30 min of infection with *Sp*, 94% of PMNs remained viable, i.e., membrane impermeable to propidium iodide (PI; Figure S2a), and PMNs killed 70% of opsonized *Sp* (Figure S2b). Infection with *Sp* triggered >7-fold increases in NETosis, PMN apoptosis, and ROS production (Figures S2c, e-f), and >2-fold increases in MMP and NE release (Figure S2d, g). (The relative log-fold changes in various activities of infected PMN parameters induced by *Sp* infection are provided in a radar plot; Figure S2h).

We next profiled the effect of HXA_3_ and fMLP on *Sp*-induced responses of infected PMNs. fMLP treatment resulted in a slight increase in membrane-permeant PMNs compared to HBSS or HXA_3_ treatment (10% versus 6%; Figure 5a). Nevertheless, fMLP- and HXA_3_-treated PMNs were equally proficient as HBSS-treated PMNs at opsonophagocytic killing (Figure 5b). The presence or absence of fMLP or HXA_3_ also did not affect NETosis or MMP secretion by infected PMNs (Figure 5c, d). fMLP treatment resulted in slightly higher levels of apoptosis, reflected by surface levels of Annexin V compared to untreated or HXA_3_-treated PMNs (Figure 5e), a finding consistent with the observation that HXA_3_ diminishes PMN apoptosis (37). Finally, HXA_3_ resulted in slightly higher ROS production than fMLP (10% versus 8%, Figure 5f).

**Figure 5.**
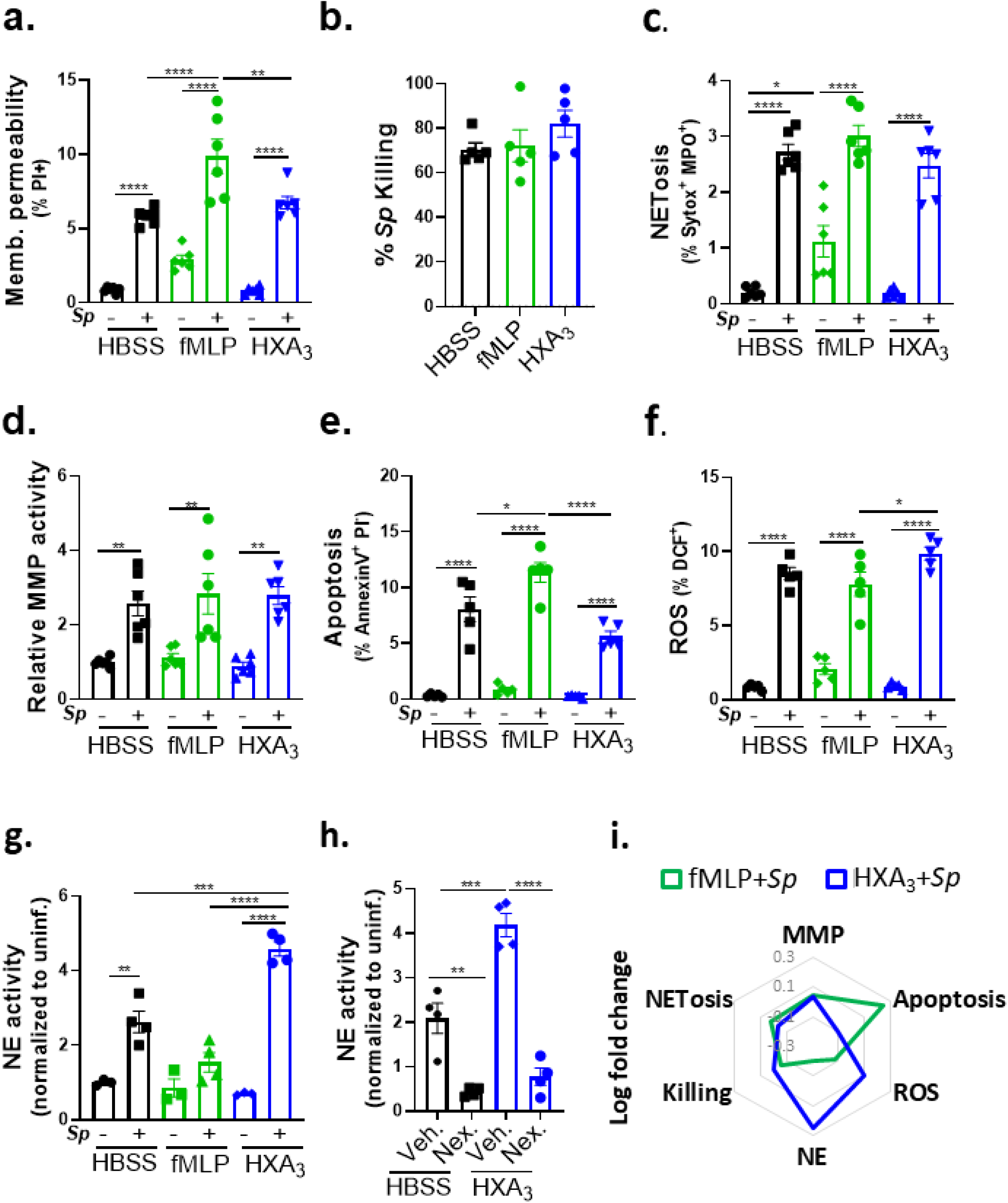
HXA_3_ enhances NE secretion by *Sp*-infected PMNs. 1 × 10^6^ PMNs were infected with 1 × 10^7^ *Sp* after treatment with control HBSS, 10 µM fMLP, or 10 nM HXA_3_ methyl ester (“HXA_3_”), and evaluated for functional performance via **(a)** PMN membrane permeability determined by propidium iodide staining (PI^+^), **(b)** opsonophagocytic killing quantitated by plating for CFU, **(c)** NETosis determined by Sytox and anti-MPO staining (Sytox^+^ MPO^+^), **(d)** released MMP activity by substrate conversion and expressed relative to uninfected PMNs, **(e)** apoptosis determined by lack of straining by propidium iodide and positive staining of Annexin V (PI^-^ Annexin V^+^), **(f)** ROS production by intracellular oxidation of substrate (DCF^+^), and **(g)** released NE activity by substrate conversion and expressed relative to uninfected PMNs. **(h)** *Sp*-infected PMNs were treated with HXA_3_ methyl ester in the presence or absence of 50 µM Nexinhib20 (Nex.) and relative NE activity in supernatant quantitated by substrate conversion as in panel g. **(i)** Radar plot summary of log fold change in PMN activities in **(a-g)**. Each panel shown is representative of three independent experiments. Error bars represent mean ± SEM. Statistical analysis was performed using ordinary one-way ANOVA: *p-value < 0.05, **p-value < 0.01, ***p-value < 0.001, ****p-value < 0.0001.

The greatest chemoattractant-dependent difference detected in Sp-infected PMNs was NE activity. fMLP stimulation appeared to diminish NE activity compared to PBS, although this difference did not reach statistical significance (Figure 5g). In contrast, HXA_3_ resulted in an almost 2-fold increase relative to the control (P<0.001). NE has been implicated in severe lung injury during *Sp* infection (49, 50) and is delivered by PMNs largely through the release of exosomes and primary granules (51). The increase in HXA_3_-triggered NE activity was eliminated by Nexinhib20, which blocks NE release by both exosomes and primary granules (52) (Figure 5h, “Nex”). The relative changes in various activities of infected PMN parameters induced by fMLP or HXA_3_ are summarized in Figure 5i.

### PLY-producing *Sp* promotes release of NE and primary granules in a 12-LOX-dependent manner during experimental lung infection

Given the enhanced NE release by HXA_3_-stimulated, *Sp*-infected PMNs *in vitro*, we assessed in vivo degranulation of primary granules, a major mechanism of NE release (51) of PMNs. 18 hours after i.t. infection of BALB/c mice with *Sp,* we measured the relative level of the primary granule marker CD63 on the surface of pulmonary PMNs. CD63 surface expression was 3.5-fold higher on PMNs from lungs of mice infected with WT *Sp* compared to uninfected mice (Figure 6b, “Uninf.- vs. “WT”). This elevated level was reduced by 25% during infection with *Sp Δply* (P<0.05; Figure 6b, “*Δply*”), suggesting that PLY-induced HXA_3_ production contributed significantly to degranulation. Consistent with this, inhibition of 12-LOX with CDC after infection with WT *Sp* infection resulted in a similar decrease in PMN CD63 surface expression (Figure 6a, “CDC”).

**Figure 6.**
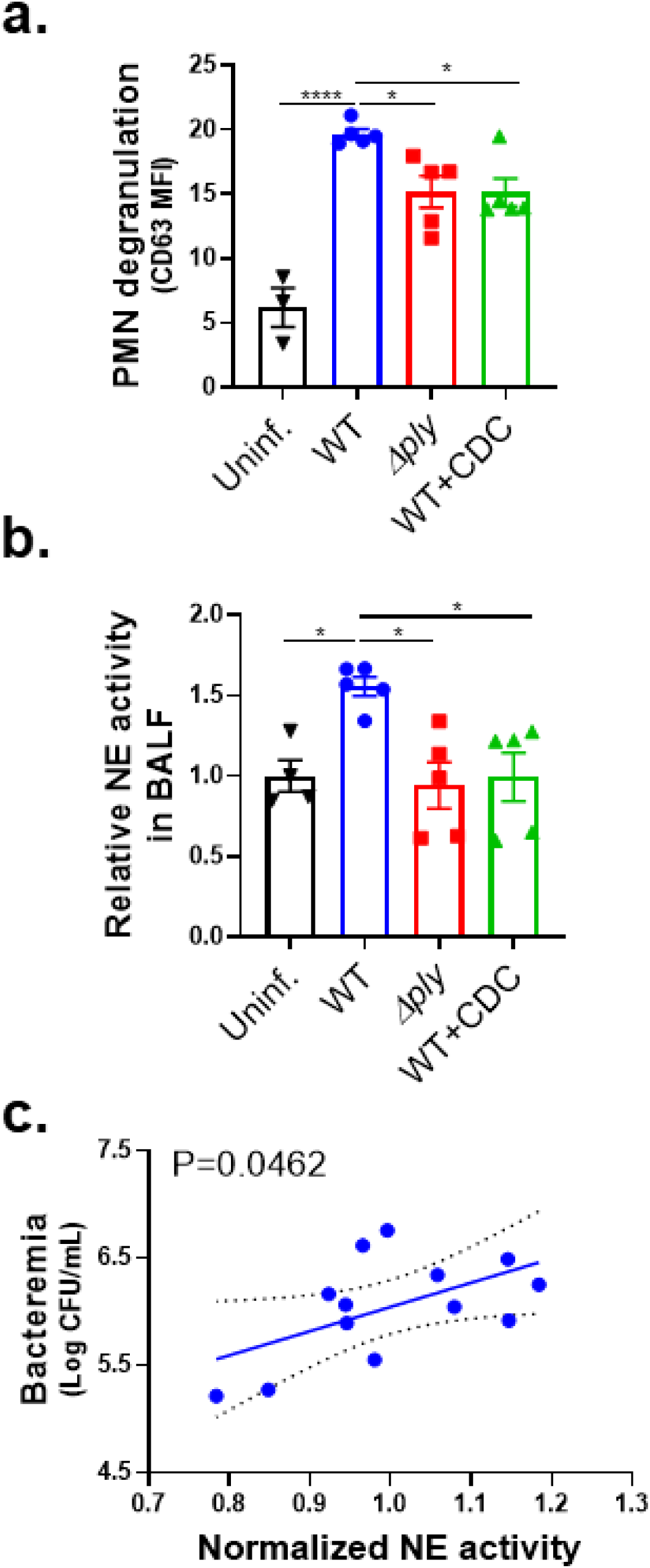
PLY-producing *Sp* promotes release of NE and primary granules in a 12-LOX-dependent manner during experimental lung infection. BALB/c mice were infected *i.t.* with 1 × 10^7^ CFU WT or *Δply Sp* for 18 h, with or without *i.p* injection of 8 mg/kg of the 12-LOX inhibitor CDC. **(a)** NE activity in cell-free BALF determined by substrate conversion, expressed relative to the NE activity in cell-free BALF from uninfected mice. **(b)** FACS analysis of degranulation determined by CD63 expression on Ly6G^+^ lung infiltrating PMNs. **(c)** Correlation between normalized NE activity in (a) and bacteremia determined by enumerating CFU in serum. Each panel shown is representative of three independent experiments, or pooled data from three independent experiments. Error bars represent mean ± SEM. Statistical analysis was performed using ordinary one-way ANOVA: *p-value < 0.05, ***p-value < 0.001, ****p-value < 0.0001.

To determine if PMN degranulation corresponded to increased pulmonary NE levels bronchial alveolar lavage fluid (BALF) of BALB/c mice at 18 h.p.i. was centrifuged to remove PMNs and other cells, and then assessed for NE activity. Activity was 50% higher in WT *Sp*-infected mice compared to uninfected mice or mice infected with *Sp Δply* (Figure 6b, “*Δply*”), a finding consistent with previous reports (7). CDC treatment of infected mice, which dramatically decreases PMN lung infiltration (29), reduced BALF NE to levels indistinguishable from that of uninfected mice (Figure 6b, “CDC”). In mice infected with WT *Sp*, BALF NE activity significantly correlated with bacteremia (Figure 6c). These findings suggest that PLY-triggered HXA_3_ promotes lung-infiltrating PMNs to release NE during pulmonary *Sp* challenge, thus enhancing bacteremia.

### Inhibition of NE release mitigates disruption of the lung epithelial barrier and bacteremia following *Sp* lung infection

NE degrades epithelial cell junctions and extracellular matrices *in vitro* (53, 54) and has been implicated in the pathogenesis of several human disorders that involve inflammatory damage (51). To determine if inhibition of PMN degranulation or NE activity protects lung barrier function during *Sp* infection in mice, we delivered the degranulation inhibitor Nexinhib20 or the NE inhibitor Sivelestat (16) by intraperitoneal (*i.p.*) injection to BALB/c mice (see Methods), followed by *Sp* lung challenge. Neither inhibitor altered bacterial lung burden or PMN infiltration at 18 h.p.i (Figure 7a-b). Nexinhib20 significantly diminished PMN degranulation, measured by surface CD63, compared to vehicle-treated mice (Figure 7c); Sivelestat did not achieve significant effect. Notably, both Nexinhib20 and Sivelestat prevented an increase in the NE activity of BALF (Figure 7d).

**Figure 7.**
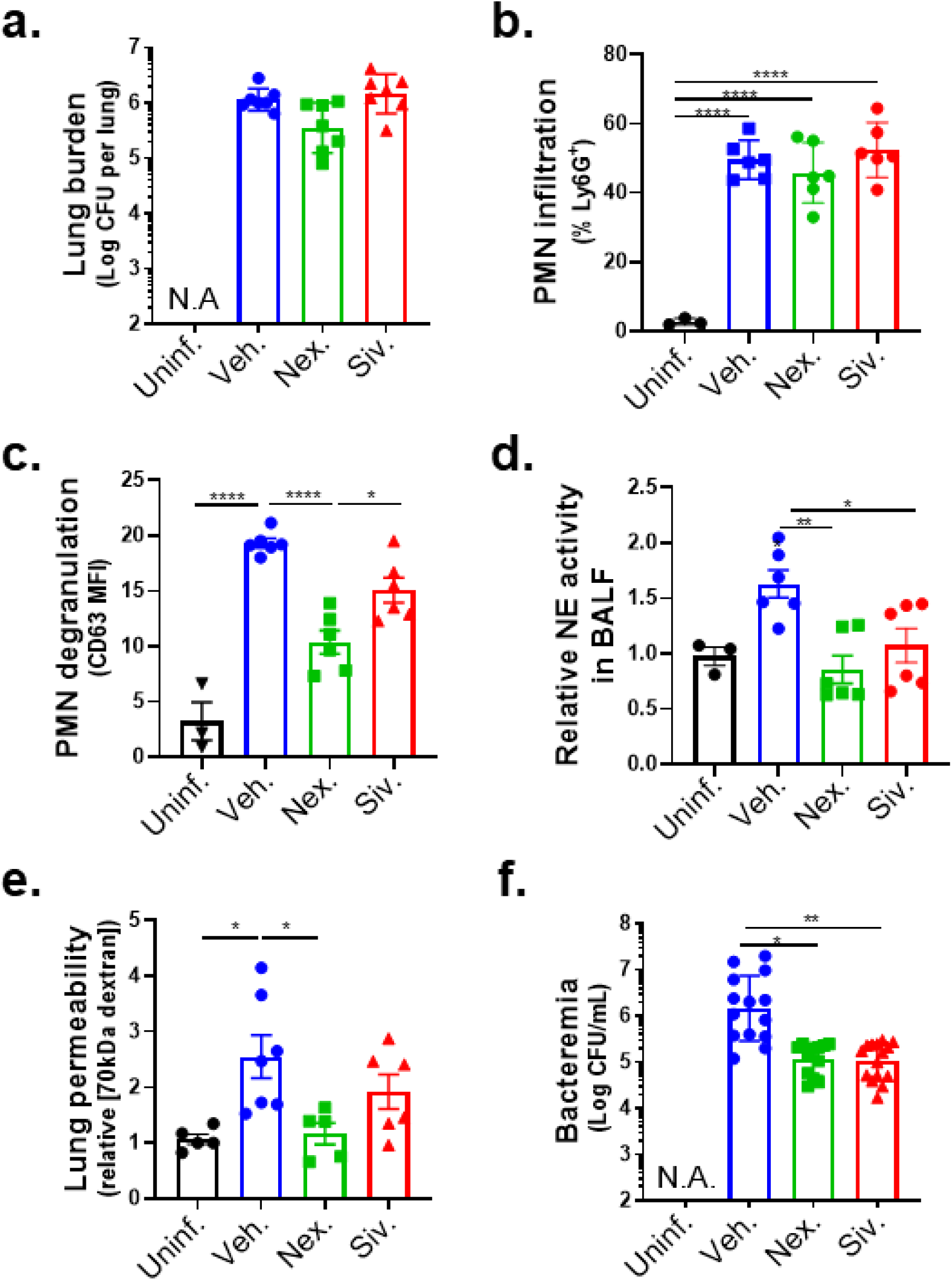
Inhibition of NE release mitigates disruption of the lung epithelial barrier and bacteremia following *Sp* lung infection. BALB/c mice were infected *i.t.* with 1 × 10^7^ CFU WT *Sp* for 18 h, with or without *i.p* injection of 30 mg/kg Nexinhib20 (Nex) or 30 mg/kg Sivelestat (Siv) one hour prior to infection. **(a)** Bacterial lung burden determined by measuring CFU in lung homogenates; **(b)** PMN infiltration determined by flow cytometric enumeration of Ly6G^+^; **(c)** Degranulation determined by CD63 expression on Ly6G^+^ lung infiltrating PMNs by FACS; **(d)** Relative NE activity in BALF determined by substrate conversion; **(e)** Lung permeability determined by measuring the concentration of 70 kD FITC-dextran in lung relative to serum after *i.v.* administration; and **(f)** Bacteremia determined by enumerating CFU in serum. Each panel is representative of three independent experiments, or pooled data from three independent experiments. Error bars represent mean ± SEM. Statistical analysis was performed using ordinary one-way ANOVA: *p-value < 0.05, **p-value < 0.01, ****p-value < 0.0001.

The decrease in PMN degranulation associated with Nexinhib20 significantly protected the lung epithelial barrier, reducing epithelial barrier permeability to intravenous 70 kDa FITC dextran by 55% (Figure 7e, “Nex.”); Sivelestat treatment exhibited a similar trend, reducing permeability by 25% (Figure 7e, “Siv.”). Importantly, both inhibitors diminished bacteremia significantly by >10-fold (Figure 7f). These data suggest that NE release by HXA_3_-activated lung infiltrating PMNs contributes to barrier disruption.

## Discussion

Lung infections by viral and bacterial pathogens, especially multi-drug resistant bacteria, remain a major cause of death and require searches for therapies that target infection-associated pathogenic host processes (4, 7). Pulmonary infiltration by PMNs can drive lung damage and concomitant transepithelial movement of pathogens, including *Sp* (55–57), leading to life-threatening systemic infection. Indeed, transepithelial migration of PMNs in response to activation of the 12-LOX pathway disrupts cultured epithelial monolayers (29) and promotes lethal bacteremia in a mouse *Sp* lung challenge model (17). However, PMNs are also key immune cells critical for early defense against *Sp* infection (58). Hence, efficacious host-directed therapies to combat *Sp* spread must selectively target PMN effector mechanisms that promote barrier disruption while leaving intact activities required for pathogen control. Identification of the critical pathologic activities of PMNs during *Sp* infection of the lung requires model systems that faithfully reflect key features of PMN-*Sp* interactions at the respiratory mucosa.

The bronchial BSC-derived ALI epithelial model recapitulates important aspects of the architecture of *bona fide* airway epithelium, including the diversity of cell types and the formation of mature apical junction complexes that facilitate a functional mucosal barrier (45, 59). Here, we show that *Sp* infection of human and murine BSC-derived ALI monolayers mirror essential features of epithelial barrier breach following pulmonary *Sp* challenge in mice (17, 60), including the requirement for PMN transmigration that is entirely dependent on 12-LOX and partially dependent on PLY (29). PLY does not trigger detectable PMN transmigration or concomitant bacterial translocation after genetic ablation of 12-LOX pathway, suggesting that PLY-triggered pro-inflammatory and barrier disrupting activity in the lung is entirely due its ability to stimulate this pathway. That a PLY-deficient *Sp* was still capable of triggering 12-LOX-dependent PMN migration across ALI monolayers, albeit at lower than wild-type levels (Figure 2), is consistent with previous work indicating that *Sp* is also capable of stimulating PMN transmigration via PLY-independent means (29).

Chemotactic cues can have remarkably diverse effects on PMNs, including altering effector functions, antimicrobial activity, and inflammatory potential (23, 61). For example, in models of sterile lung injury, infiltrating PMNs are apoptotic and produce tissue-repair molecules such as TGF-β, VEGF, and resolvins (62–64). Conversely, in cystic fibrosis (CF) models, PMNs undergo transcriptional changes that reduce bactericidal activity and enhance tissue-damaging degranulation (40, 65). Similarly, PMNs that migrate into COVID-19-infected airways display a hyperinflammatory phenotype that drives lung pathology (41). Here, we show that chemotactic cues ultimately lead to divergent infection outcomes in *Sp* infection of ALI monolayers. Based on analogy to mucosal infection by other pathogens (32–34), HXA_3_ was previously deemed likely to be the 12-LOX-dependent PMN chemoattractant driving acute inflammation during *Sp* infection (17, 18). Here, the experimental utility of 12-LOX-deficient ALI monolayers permitted the definitive identification of HXA_3_as indeed being sufficient to induce PMN transmigration and mucosal barrier disruption triggered by *Sp* infection. In turn, this finding was essential to permit a direct comparison of *Sp*-driven chemotaxis with that triggered by a well-studied control chemoattractant, fMLP (23, 66), revealing that HXA_3_- promoted specific pro-inflammatory conditioning of PMNs is critical for epithelial monolayer destruction.

The identification of HXA_3_ as sufficient for PMN-mediated mucosal barrier breach during infection by *Sp* prompted an exploration of pathologically important HXA_3_ responses. Changes in PMN physiology upon stimulation by purified HXA_3_ include increased calcium flux, NETosis, and anti-apoptotic programs (32, 37, 38), but here we investigated HXA_3_ response in the context of *Sp* infection. By far the largest difference upon *ex vivo* treatment of *Sp*-infected PMNs with HXA_3_ compared to fMLP was a 4-fold higher level of NE activity (Figure 5). HXA_3_ alone does not enhance PMN NE activity (Figure 5), indicating that this response requires co-stimulation by both bacteria and chemoattractant and emphasizing the importance of including microbial agents in studies of PMN responses to infection-triggered chemotactic agents. Moreover, pulmonary PMNs from mice challenged *i.t.* with *Sp* exhibited PLY- and 12-LOX-promoted elevation of degranulation, a means to release NE, as well as elevated NE levels in BALF, indicating that HXA_3_ triggered NE release during mouse lung infection as well (Figure 6).

Disease manifestation in response to pathogens can be mitigated either by effective actions of the host immune system to clear the microbe or by control of infection-triggered immune responses that are detrimental to the host (67). NE, along with other serine proteases, contributes to *Sp* killing by PMNs *ex vivo* (10). However, we found that inhibition of NE activity during mouse lung infection by pretreatment with the NE inhibitor Sivelestat did not affect bacterial lung burden (Figure 7), nor did it alter PMN lung infiltration. Rather, inhibition of NE, which degrades extracellular matrix components (68) and alveolar epithelial cell junction proteins (69) that maintain epithelial integrity (70), decreased bacteremia by more than 90%. These findings indicate that, in the mouse model, the pathological activities of NE outweigh any beneficial role in direct pathogen killing (10, 12).

Nexinhib20 blocks formation of exosomes and degranulation of primary granules (52), the two means by which NE is released from PMNs. During mouse infection, this inhibitor diminished degranulation of lung PMNs as well as NE activity in BALF. Although Nexinhib20 diminishes surface localization of adhesion molecules and can limit PMN recruitment to sites of tissue damage (71), we found that this inhibitor did not alter PMN infiltration into the lung post-*Sp* challenge. Nexinhib20 has been shown to ameliorate PMN-directed tissue damage in models of myocardial ischemia-reperfusion (71) and pulmonary LPS-induced injury (72). Here we demonstrated the ability of the drug to mitigate injury during microbial infection. Treatment with Nexinhib20, like treatment with Sivelestat, did not alter pulmonary bacterial load (Figure 7), yet bacteremia was decreased >10-fold, corresponding to protection of pulmonary barrier function measured by leakage of 70 kDa dextran (Figure 7). Primary granules contain numerous proteases that may have diverse physiologically activities (73, 74), such as the activation or inactivation of cytokines and other biologically active host factors (75, 76), that may impact the course of *Sp* infection, so further characterization of the effects of Sivelestat and Nexinhib20 *in vivo* is required to garner a full understanding of how they diminish bacteremia.

NE has been implicated in the pathogenesis of several human disorders that involve inflammatory damage, including CF, chronic obstructive pulmonary disease, bronchopulmonary dysplasia, and acute respiratory distress syndrome (ARDS) (51). The pathogenic role of NE activity during *Sp* infection of the mouse lung revealed here is likely reflected in human infection because higher NE levels in BALF and serum of patients with bacterial pneumonia is associated with worse clinical outcomes (77–79). Sivelestat is clinically approved for the treatment of ARDS in Korea and Japan (80) and for COVID-19 induced ARDS in China (81). Although studies of efficacy in patients have yielded inconsistent results (51, 82–84), ongoing efforts to improve delivery, e.g., by nebulizer, have yielded favorable results in improving efficacy and limiting adverse effects (85). Similarly, intrapulmonary delivery of Nexinhib20-loaded nanoparticles in experimental animals increases drug availability and decreases LPS-induced acute lung injury (72). Future studies are required to determine the efficacy of NE inhibition in limiting *Sp* systemic disease.

Finally, HXA_3_ production is a conserved mucosal inflammatory response in a multitude of bacterial infections, and possibly in acute lung injury, asthma, and inflammatory bowel syndrome (22, 31, 32, 34, 86). Given the prominent role of PMNs in mediating tissue damage, targeted mitigation of HXA_3_-triggered changes in PMNs that promote damage but do not compromise host defense has potential efficacy for a broad range of disorders. The identification of such changes, such as excessive NE release, is an important step in developing such strategies.

## Materials and Methods

### Bacterial strains and growth conditions

Mid-exponential growth phase aliquots of *S. pneumoniae* TIGR4 (serotype 4) were grown in Todd-Hewitt broth (BD Biosciences) supplemented with 0.5% yeast extract in 5% CO_2_ and Oxyrase (Oxyrase, Mansfield, OH), and frozen in growth media with 20% (v/v) glycerol. Bacterial titers in aliquots were confirmed by plating serial dilutions on Tryptic Soy Agar plates supplemented with 5% sheep blood (blood agar) (Northeast Laboratory Services, Winslow, ME). The TIGR4 PLY-deficient mutant (*Δply*) was a gift from Dr. Andrew Camilli (Tufts University School of Medicine, MA). For experiments, *S. pneumoniae* strains were grown in Todd-Hewitt broth, supplemented with 0.5% yeast extract and Oxyrase, in 5% CO_2_ at 37°C and used at mid-log to late log phase.

### Murine infections

BALB/c mice, C57BL/6J mice, and *Alox15* knockout (*Alox15*^−/−^) mice (B6.129S2- *Alox15*tm1Fun/J) were obtained from Jackson Laboratories. All animal experiments were performed in accordance with Tufts University Animal Care and Use Committee approved protocols. BALB/c mice were intratracheally challenged with 1×10^7^ colony forming units (CFU) of *S. pneumoniae* in 50 μl phosphate-buffered saline (PBS) to induce pneumococcal pneumonia. Control mice received PBS. The role of 12-LOX on *S. pneumoniae*-induced inflammation in BALB/c mice was investigated by inhibiting this enzyme with cinnamyl-3,4-dihydroxy-α-cyanocinnamate (CDC) at 8 mg/kg, in 3% DMSO, 3% cremaphor EL (CrEL) in PBS as the vehicle. CDC was injected intraperitoneally (*i.p.*) twice daily, starting one day before infection. The role of NE on *S. pneumoniae*-induced inflammation was studied in BALB/c mice by treatment with Nexinhib20, which blocks release of primary granules (52) or Sivelestate, which inhibits this enzyme (16, 87). A single dose of Nexinhib20 at 30 mg/kg, in 3% DMSO, 3% CrEL in PBS, or Sivelestat at 30 mg/kg in PBS was injected *i.p.* 1 hour prior to infection. Mice were euthanized at 18 h.p.i.. Blood was obtained by cardiac puncture. Bronchoalveolar lavage fluid (BALF) was collected by washing the lungs twice with 1 ml PBS via a cannula, then whole lungs were removed and bacterial burden enumerated by plating lung homogenate on blood agar plates.

### Assessing lung barrier function

For assessment of lung permeability, mice were intravenously injected with 70 kDa MW FITC-Dextran at 5 mg/kg 30 minutes prior to euthanasia. Whole lungs were isolated and homogenized in 1 ml PBS, which was then subjected to fluorescence quantitation using a Synergy H1 plate reader (BioTek). Readouts were normalized to fluorescence in the serum of the same animal, diluted 1:10 in PBS.

### Measuring PMN infiltration and degranulation

For flow cytometric quantitation of lung PMNs, mice were euthanized at 18 h.p.i. and lung tissues were digested with 1 mg/ml Type II collagenase (Worthington) and 50 U/ml Dnase (Worthington) to obtain a single-cell suspension. Cells present in the suspension were stained on ice for 30 minutes with APC-conjugated anti-Ly-6G (clone 1A8) or PE-conjugated anti-CD63 (Biolegend) and then washed two times in FACS buffer (Biolegend). Cells were analyzed using a FACSCalibur flow cytometer (BD Biosciences) and the fluorescence intensities of the stained cells were determined. Collected data were analyzed using FlowJo software (v10.7, BD) to determine the numbers of infiltrating (Ly6G^+^) PMNs, and their level of degranulation (mean fluorescence intensity of CD63).

### Establishment of epithelial air-liquid interface monolayers

Human bronchial basal cells were isolated and expanded from lung tissue harvested from donors without lung disease through the New England Organ Bank under an IRB-approved protocol (MGH #2010P001354). In brief, using a previously published basal cell isolation protocol (45, 88), EpCAM^+^ epithelial basal cells were isolated from human trachea and mainstem bronchi tissue. Mouse airway basal cells were obtained from C57BL/6J (B6) or *Alox15*^−/−^ mouse trachea.

Harvested basal cells were cultured in complete small airway epithelial growth media (SAGM) (Lonza, Cat. CC-3118), with propagation for up to 10 passages, using the dual SMAD inhibition protocol (45).To generate monolayers permissive to modeling PMN transmigration (59), Transwells with permeable (3 µm pore size) polycarbonate membrane inserts and a culture area of 0.33 cm^2^ (Corning product #3415) were collagen coated and seeded with 80 µl of the airway basal cells suspension (containing > 200,000 cells) in SAGM, resulting in a density of >6000 cells/mm^2^, and submerged in complete SAGM for airway basal cell recovery and expansion for 1–2 days to ensure monolayer confluence. The media in both chambers was then replaced with complete Pneumacult-ALI medium (StemCell Technology, Cat. 05001) for an additional day. To initiate air-liquid interface, ALI medium in the chamber contacting the cell apical surface was removed (designated as day 0). Media was changed every 1-2 days during differentiation.

ALI monolayers used in experiments were cultured for at least 21 days to allow for full maturation of both cilia and goblet cells, but no more than 34 days to avoid overgrowth or loss of epithelial barrier (42). Transepithelial electrical resistance was assessed using a voltmeter (EVOM2, Epithelial Voltohmmeter, World Precision Instruments, Inc.) prior to migration assays to ensure the establishment of a polarized epithelial barrier.

### Infection of ALI monolayers

*S. pneumoniae* grown to log phase was washed and resuspended to 5×10^8^ CFU/ml in Hanks’ balanced salt solution (HBSS) supplemented with 1.2 mM Ca^2+^ and 0.5 mM Mg^2+^. 25 μl of bacterial suspension was added to the apical surface of the ALI monolayers (grown on the underside of the Transwells) by inverting the Transwells and incubating at 37°C with 5% CO_2_ for 2 hours to allow for attachment and infection of the ALI monolayers. After treatment, Transwells were placed in 24-well receiving plates containing HBSS with Ca^2+^ and Mg^2+^, and to allow for bacteria translocation, incubated for an additional 2 hours with or without the addition of 1×10^6^ PMNs to the basolateral chamber. Detection of basally added horseradish peroxidase (HRP) in the apical chamber was used to assess ALI monolayer barrier integrity post-treatment. Buffer in the basolateral chambers was sampled and bacterial translocation across ALI monolayers was evaluated by plating serial dilutions on blood agar plates. Bacterial migration index was calculated as total CFUs in the basolateral chamber normalized to infection inoculum.

### Production of cell supernatants containing HXA_3_

Epithelial cell supernatants were generated from B6 ALI monolayers infected with 1×10^7^ WT or *Δply S. pneumoniae* for 1 hour at 37°C with 5% CO_2_, and then placed in 24-well receiving plates containing HBSS with Ca^2+^ and Mg^2+^ in the apical chamber for an additional 2 hours to allow for HXA_3_ generation. The apical chamber supernatants were then collected and transferred to new Transwells with ALI monolayers for PMN transmigration assays.

### PMN transepithelial migration assays

Whole blood obtained from healthy human volunteers under an IRB-approved protocol (Tufts University protocol #10489) was used to isolate neutrophils using the Easysep direct human neutrophil isolation kit (Stemcell), and 1×10^6^ PMNs were added to the basolateral chamber after two hours of apical infection of the ALI monolayers with *S. pneumoniae*. Purified HXA_3_ methyl ester (Cayman) at 10 nM and fMLP (Sigma) at 10 μM were supplemented apically as indicated. To test the effect of HXA_3_-containing cell supernatants, the apical media was replaced with cell supernatants prepared as described above.

After two hours of transmigration, PMNs in the apical chamber were quantified by MPO activity assay, as described (29). Briefly, 50 μl of 10% Triton X-100 and 50 μl of 1 M citrate buffer were added to lyse transmigrated PMN, and 100 μl of lysed PMNs from each well was transferred to a 96-well plate. 100 μl of freshly prepared 2,2’-azinobis-3-ethylbenzotiazoline-6-sulfonic acid (ABTS) with hydrogen peroxide solution was added to each well and incubated in the dark at room temperature for 5-10 minutes. Absorbance at a wavelength of 405 nm was read on a microplate reader and measurement was converted to neutrophil number using a standard curve.

### Fluorescence microscopy assessment of ALI monolayer integrity

At the end of the two hours of infection, followed by two hours of PMN transmigration across ALI monolayers, the degree of cell confluency of ALI monolayers on Transwell filters was assessed by fluorescence microscopy. To prepare samples for fluorescence microscopy, 4% paraformaldehyde fixed ALI monolayers were permeabilized with 0.1% Triton-X 100 in PBS with 3% BSA. ALI monolayers were then stained with DAPI (for nuclei) and Alexa Fluor 594 phalloidin (for F-actin), and visualized on excised filters with a Leica SP8 spectral confocal microscope (Leica). Epithelial cell retention on filters was quantitated by counting of DAPI-stained epithelial cell nuclei per field of view, carried out with CellProfiler pipeline optimized with size and roundness exclusion criteria for epithelial cell nuclei identification. Counts were normalized to uninfected controls.

### Neutrophil elastase and metalloprotease activity

NE activity and MMP activity in soluble fraction of BALF from infected mice or PMN supernatants from 1×10^6^ PMNs challenged with 1×10^7^ CFU *S. pneumoniae* was determined using a PMN Elastase Fluorometric Activity Assay Kit (Abcam) and Fluorogenic MMP Substrate (Mca-PLAQAV-Dpa-RSSSR-NH2) (R&D Systems), respectively, following manufacturer’s instructions. The area under the curve of kinetic substrate conversion curves over two hours was measured with a Synergy H1 plate reader (BioTek) and normalized to uninfected controls.

### Opsonophagocytic (OPH) killing

The ability of neutrophils to kill pneumococci was assessed *ex vivo* as described previously (89), with modification. Briefly, 1×10^6^ PMNs were incubated with 5 x10^3^ *S. pneumoniae* grown to mid-log phase and pre-opsonized with 10 μl rabbit complement (Pel-Freez) in 100 μl reactions in HBSS with Ca^2+^ and Mg^2+^. The reactions were incubated for 45 minutes at 37°C. Percent killing in comparison to incubations with no PMNs was determined by plating serial dilutions on blood agar plates.

### Reactive oxygen species (ROS) production

Neutrophils were resuspended in HBSS with Ca^2+^ and Mg^2+^ containing 10 μM 2’,7’- dichlorodihydrofluorescein diacetate (DCF) (Molecular Probes) to a final concentration of 1×10^7^ cells/ml and gently agitated for 10 minutes at room temperature. 1×10^6^ DCF-containing neutrophils were challenged with *Sp* at an MOI of 10 for 30 minutes at 37°C, then washed and resuspended in FACS buffer for analysis by a FACSCalibur flow cytometer (BD Biosciences). Collected data were analyzed using FlowJo software (v10.7, BD) to determine the numbers of ROS-producing DCF-positive cells.

### Neutrophil extracellular trap formation (NETosis) and apoptosis by flow

1×10^6^ neutrophils were challenged with *Sp* at an MOI of 10 for 30 minutes at 37°C, then washed and resuspended in FACS buffer. For NETosis analysis, cells were stained with a plasma membrane-impermeable DNA-binding dye, SYTOX™ AADvanced™ (Life Technologies, Carlsbad, CA), rabbit anti-myeloperoxidase (Abcam ab45977), and secondary goat anti-rabbit-Alexa Fluor 568 antibody (Invitrogen). For apoptosis analysis, cells were stained with FITC-conjugated Annexin V (BioLegend, San Diego, CA, USA), and propidium iodide (PI). Samples were read on a FACSCalibur flow cytometer (BD Biosciences), and collected data were analyzed using FlowJo software (v10.7, BD) to determine percent NETosis (MPO^+^ SYTOX^+^), and percent apoptosis (Annexin^+^ PI^-^).

### Presentation of data and statistical analyses

Statistical and correlation analysis was carried out using GraphPad Prism (GraphPad Software, San Diego, CA). p values <0.05 were considered significant in all cases. For bacterial burdens, geometric mean ± geometric SD is shown; for all other graphs, the mean values ± SEM are shown. Due to intrinsic donor-to-donor variability of human PMN transmigration efficacy, experiments involving human donors were normalized within each experiment before pooling individual experiments. The conclusions drawn were those found to be reproducible and statistically significant across independent experiments.

## Acknowledgements

We thank Andrew Camilli for strains and Elsa Bou Ghanem, Stephania Libreros, Amanda Pulsifer, Rodney K. Tweten, Byran P. Hurley, and Beth A. McCormick for protocols and helpful discussions. This work was supported by NIH Awards AG071268 to JL and JM, and NIH Awards AI15081 and AI152499 to JMV.

## Supplemental Figure Legends

**Supplemental Figure 1. PM**Ns **are required for epithelial cell detachment and barrier breach.** Human BSC-derived ALI monolayers were apically infected with 1 × 10^7^ WT or *Δply Sp* without basolateral PMNs. **(a)** After two hours, monolayer integrity was assessed by fluorescence confocal microscopy after staining nuclei with DAPI and F-actin with fluorescent phalloidin. For clarity, images shown are of extended projections (all z-sections collapsed into 1 plane). Scale bar = 40 μm for all images. **(b)** Epithelial retention was quantitated by enumerating epithelial cell nuclei relative to uninfected ALI. **(c)** Epithelial permeability was measured by HRP flux relative to monolayers infected with WT *Sp*. **(d)** *Sp* translocation was quantitated by measuring basolateral CFU. Each panel is a representative of three independent experiments. Error bars represent mean ± SEM.

**Supplemental Figure 2.** S***p* infection alters PMN functional response profile.** 1 × 10^6^ PMNs were uninfected or infected with 1 × 10^7^ *Sp* and evaluated for functional performance via **(a)** PMN membrane permeability determined by propidium iodide staining (PI^+^), **(b)** opsonophagocytic killing with or without the addition of complement opsonin, quantitated by plating for CFU, **(c)** NETosis determined by Sytox and anti-MPO staining (Sytox^+^ MPO^+^), **(d)** released MMP activity by substrate conversion and expressed relative to uninfected PMNs, **(e)** apoptosis determined by lack of straining by propidium iodide and positive staining of Annexin V (PI^-^ Annexin V^+^), **(f)** ROS production by intracellular oxidation of substrate (DCF^+^), or **(g)** released NE activity by substrate conversion and expressed relative to uninfected PMNs (See Materials and Methods). **(h)** Radar plot summary of log fold change in PMN functional performance in **(c-g)**. Each panel shown is representative of three independent experiments. Error bars represent mean ± SEM. Statistical analysis was performed using ordinary one-way ANOVA: **p-value < 0.01, ***p-value < 0.001, ****p-value < 0.0001.

